# STAR+WASP reduces reference bias in the allele-specific mapping of RNA-seq reads

**DOI:** 10.1101/2024.01.21.576391

**Authors:** Rebecca Asiimwe, Dobin Alexander

## Abstract

**Summary:** Allele-specific expression (ASE) is an important genetic phenomenon that impacts an individual’s phenotype and is relevant in various biological and medical contexts. Next-generation RNA sequencing technologies provide an unprecedented opportunity to measure ASE genome-wide across all heterozygous alleles expressed in a given sample. One of the major obstacles to the accurate calculation of ASE from RNA-seq data is the reference mapping bias, i.e., the preferential misalignment of the reads to the reference allele. Here, we present STAR+WASP, our reimplementation of WASP, a highly accurate algorithm for reducing the reference bias (Van De Geijn *et al*. 2015). We show that STAR+WASP is an order of magnitude faster than WASP while significantly reducing reference bias and providing ASE estimations similar to the original WASP algorithm.

**Availability and Implementation:** STAR+WASP is implemented within STAR as an integrated C++ module. STAR+WASP is open-source software, freely accessible at: http://code.google.com/p/rna-star/.

**Supplementary information:** Supplementary data are available at Bioinformatics online.

## Introduction

Allele-specific expression (ASE) is a genetic phenomenon where the two alleles of a particular gene are not expressed equally in an individual’s cells. The importance of allele-specific expression lies in its potential to influence an individual’s phenotype and its relevance in various biological and medical contexts, such as genetic diversity and evolution, gene regulation, disease susceptibility, pharmacogenetics, etc. Next-generation RNA sequencing technologies provide an unprecedented opportunity to measure ASE genome-wide across all heterozygous alleles expressed in a given sample. Calculating allele-specific expression in RNA-seq data involves mapping reads to the reference genome, identifying reads that overlap heterozygous variants, assigning the reads to one of the alleles, and calculating allelic count imbalance (Castel *et al*. 2020). This approach was successfully utilized for multiple ASE-based analyses, such as identifying cis-regulated gene expression variations, nonsense-mediated mRNA decay, rare regulatory variation, and imprinting (Castel *et al*. 2015; Cosentino, Brink, and Siegel 2021).

One of the major obstacles to the accurate calculation of ASE from RNA-seq data is the mapping bias. Conventionally, quantification of ASE from RNA-seq data begins with the alignment of sequence reads to a linear reference genome, where each genomic position constitutes only the reference allele. This often leads to reference bias, i.e. preferential alignment to the reference allele and incorrect alignment of reads containing non-reference alleles (Degner *et al*. 2009; Stevenson, Coolon, and Wittkopp 2013). Although methods such as the use of consensus genomes (Kaminow *et al*. 2022), personalized genomes (Rozowsky *et al*. 2011; Groza *et al*. 2020), variant/SNP-aware aligners (Hach *et al*. 2014), graph-based aligners (Paten *et al*. 2017; Garg *et al*. 2018; Kim *et al*. 2019; Rakocevic *et al*. 2019; Rautiainen and Marschall 2020), multiple population reference genomes (Chen *et al*. 2021), and de novo assembly (Zerbino and Birney 2008), have been developed and applied to reduce reference bias, this challenge remains a substantial impediment in ASE studies.

To mitigate this problem, van de Geijn et al. (Van De Geijn *et al*. 2015) developed WASP (Workflow for Allele-Specific Processing), an allele-specific analysis method that eliminates reads with potential reference bias. WASP was shown to provide a highly accurate estimate of the ASE; however, it significantly increased computational costs. In this note, we present STAR+WASP, our re-implementation of the WASP algorithm built directly into the RNA-seq aligner STAR. We show that STAR+WASP is an order of magnitude faster than WASP while performing very similarly to the original WASP algorithm in estimating ASE and reducing the reference bias.

## Algorithm

The WASP algorithm (Van De Geijn *et al*. 2015) is based on a simple but highly effective idea **(Figure 1a)**. First, all RNA-seq reads are mapped to the reference genome, and uniquely mapped reads overlapping heterozygous single-nucleotide variants are identified. The variant alleles are flipped for each of the variants in these reads: if a read sequence contains a reference (REF) base, it is switched to the alternative (ALT) base, and if a read contains an ALT base, it is switched to the REF base. Then, the modified sequence is remapped to the reference genome. If the modified read fails to map or maps to a different locus or multiple loci, the read is flagged as reference-biased and is excluded from the ASE calculations. The WASP method significantly reduced the reference bias in simulations and real data (Van De Geijn *et al*. 2015). However, these improvements have come at the cost of substantially increased processing time and computational pipeline complexity. The main bottleneck of the WASP’s original implementation is its multistep nature, which requires writing and reading BAM files twice. To mitigate this issue, we re-implemented the WASP algorithm inside our RNA-seq aligner STAR. We follow the WASP idea of remapping the reads with a flipped genotype; however, this is done on the fly immediately after mapping the original read sequence. This avoids the input/output bottlenecks and significantly increases the processing speed. STAR+WASP attaches the “vW” tag to the alignments in the BAM output file, which indicate whether alignments pass the WASP filtering, or the reason why they are rejected (**Supplementary Table 1**).

**Figure 1.**
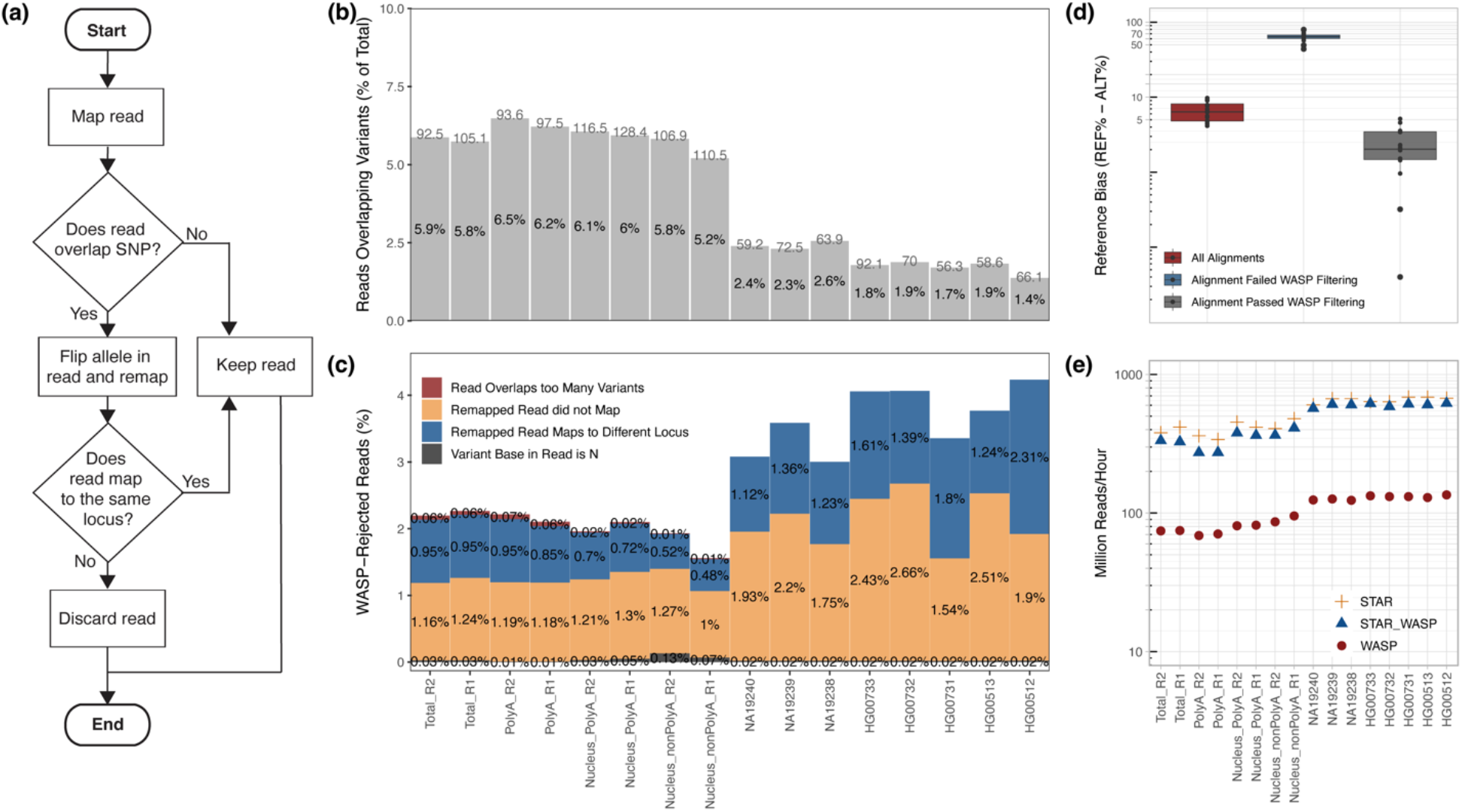
**(a)** WASP’s filtering workflow which was re-implemented in the STAR+WASP tool to reduce reference bias. Adopted from Bryce van de Geijn et al. (Van De Geijn *et al*. 2015). **(b)** Percentage of reads overlapping variants in each sample (read length is shown above each bar per sample). **(c)** Percentages of reads rejected by STAR+WASP filtering (out of reads overlapping variants) for each rejection cause. **(d)** Reference bias (REF-ALT) for unfiltered and filtered reads. **(e)** Mapping speed for STAR (no filtering), original WASP (with STAR as the mapper), and STAR+WASP algorithm.

## Benchmarks

Publicly available human RNA sequencing (RNA-seq) datasets were downloaded from the 1000 genomes (The 1000 Genomes Project Consortium *et al*. 2015) and ENCODE (The ENCODE (ENCyclopedia of DNA Elements) Project 2004) projects **(Supplementary Figure 1)**. The 16 samples included in this study represent various long RNA populations: polyA+, polyA-, and total RNA. The reads were aligned to the human reference genome (GRCh38 with Gencode v39 annotations) using STAR. The reads overlapping personal variants for each sample were filtered with WASP and STAR+WASP algorithms (see Supp. Methods).

First, we compared the reads that were deemed reference-biased and filtered out by WASP and STAR+WASP **(Supplementary Table 2, Supplementary Figures 2**,**3)**. Since the STAR+WASP algorithm was designed to resemble the WASP algorithm closely, STAR+WASP and WASP filtering results are very similar. The insignificant differences can be attributed to slight variations in filtering rules (e.g., the maximum allowed number of variants per read).

Next, we evaluated STAR+WASP performance in alleviating allelic bias. Overall, all samples **(Figure 1b)** had small but sizable proportions (1.4-6.5%) of reads that overlapped variants (these reads are labeled with a “vW” tag in the STAR+WASP BAM output). As expected, samples with longer read lengths (>90 million reads) had a higher percentage of reads overlapping variants. **Figure 1c** shows the proportions of reads that were deemed reference-biased and thus were rejected by WASP filtering for various reasons (labeled with different “vW” tag values ≠1). The main reason for rejection was the inability to remap reads after flipping the allelic variant, followed by those reads that were re-mapped to a different locus after variant flipping **(Figure 1c)**. Samples with shorter read lengths have higher rejection rates because shorter sequences are typically harder to align with high fidelity.

We further assessed the extent of reference bias introduced by aligning reads to a conventional linear reference (GRCh38) and evaluated the performance of STAR+WASP in reducing this bias. Since ground truth is unknown in real RNA-seq data, we use a proxy metric REF-ALT, defined as the difference between proportions of reads aligning to the reference and alternative alleles, averaged over all variants. Note that such differences represent allelic imbalances for each individual variant, but after averaging over all variants, the REF-ALT is expected to be close to zero. Without STAR+WASP filtering, the REF-ALT varies between 4% and 10% (median = 6.36%) for different samples **(Figure 1d, Supplementary Figure 4)**. Reads rejected by STAR+WASP filtering have large REF-ALT (44-80%) (median = 64.19%), confirming that the filtering algorithm rejects reads with strong reference bias. Alignments that passed WASP filtering have substantially smaller REF-ALT (0-5%, media = 2.03%), demonstrating that the STAR+WASP algorithm can significantly reduce the reference bias.

To demonstrate STAR+WASP computational efficiency, we benchmarked STAR, WASP, and STAR+WASP mapping speed and random access memory (RAM) usage by computing the overall runtime and RAM usage for each mapping job run with a different number of threads **(Figure 1e, Supplementary Figure 5)** on a dedicated server. STAR+WASP alignments were considerably faster (6.5 to 10.5 times) than WASP. While STAR+WASP and WASP both use STAR for the read alignment to the genome, the on-the-fly implementation of the WASP algorithm in STAR+WASP allows for much faster re-mapping and filtering of the reads. The peak memory usage of STAR+WASP was only insignificantly higher (<0.2%) than that of STAR or WASP algorithms **(Supplementary Figure 6)**. We also ran these benchmarks for all samples and tools on a shared computer cluster, which resembles more closely real-life applications, and obtained very similar results **(Supplementary Figures 7 and 8)**.

## Summary

Reference mapping bias is one of the major sources of errors in the estimation of genome-wide ASE from high-throughput RNA-seq data. We developed STAR+WASP, a re-implementation of the WASP reference bias reduction algorithm. We demonstrated that STAR+WASP produces results very similar to the original WASP tool and significantly reduces the reference bias. Since STAR+WASP is implemented within the STAR aligner codebase, it performs the filtering of reference-biased reads on the fly, allowing to decomplexify the computational pipeline, eliminate input/output bottlenecks, and increase the processing speed by an order of magnitude.

## Acknowledgments

We are grateful to Bryce van de Geijn, Graham McVicker, and Jonathan Pritchard for their fruitful discussions about the WASP algorithm.

This work was supported by the National Human Genome Research Institute of the National Institutes of Health under award number R01HG009318. The content is solely the responsibility of the authors and does not necessarily represent the official views of the National Institutes of Health.

