## Supplementary Methods for "STAR+WASP reduces reference bias in the allele-specific mapping of RNA-seq reads"

Publicly available Single-cell RNA sequencing (scRNA-seq) datasets from 16 human samples (HG00512, HG00513, HG00731, HG00732, HG00733, NA12878 Nucleus nonPolyA, NA12878 Nucleus nonPolyA Rep, NA12878 Nucleus PolyA, NA12878 Nucleus PolyA Rep, NA12878 PolyA, NA12878 PolyA Rep, NA12878 Total, NA12878 Total Rep, NA19238, NA19239 and NA19240) were downloaded from the 1000 genomes project (The 1000 Genomes Project Consortium *et al.* 2015) in form of FASTQ files, to constitute polyA+, polyA-, and total RNA populations, with reads ranging from 56x10^6^ to 128x10^6^ base pairs long - across all samples **(Supplementary Figure 1, Supplementary Table 2)**. For each sample, reads were aligned to the human reference genome (GRCh38) using STAR, STAR+WASP and WASP at 8, 16 and 32 threads per alignment. Preceding all alignments, reference genome indexes required for the alignment of all sequencing reads were generated. We supplied the reference sequences (FASTA files) of the GRCh38 genome release with the corresponding annotation file from Ensemble (gencode.v39.primary assembly.annotation.gtf). Resulting indexes were saved to disk and re-used for mapping each sample to the same reference genome. The snippet below shows how genome indexes were built.

**Building Genome Indexes**

STAR --runThreadN 16 --runMode genomeGenerate --genomeDir $genome directory --genomeFastaFiles GRCh38.primary assembly.genome.fa --sjdbGTFfile gencode.v39.primary assembly.annotation.gtf --sjdbOverhang 100

In addition to specifying FASTA files with the genome reference sequences (--genomeFastaFiles GRCh38.primary assembly.genome.fa) and annotated transcripts (--sjdbGTFfile gencode.v39.primary assembly.annotation.gtf), the number of threads (--runThreadN) used for genome building (n = 16), the genome directory (--genomeDir) to which genome indexes were to be stored for access during read alignment, and the length of the genomic sequence around the annotated junction to be used in constructing STAR’s splice junctions database (--sjdbOverhang 100) were specified. The --runMode argument for which “genomeGenerate” was specified directs STAR to run its genome indices generation module (Dobin and Gingeras 2015; Alexander Dobin 2019).

To run all STAR read mapping jobs, the number of threads to be used for read alignment was specified, in which case, each sample was run at 8, 16 and 32 threads. The path to the previously generated genome indices, read 1, and read 2 FASTQ files containing raw sequences for end-to-end alignment (--alignEndsType) were specified. As all runs were conducted in a non-shared computing environment where individual jobs utilized resources (RAM and CPU) at full server capacity, we also controlled how the genome was loaded in memory by specifying “NoSharedMemory” via the --genomeLoad argument. Importantly, to control for multimappers (reads that map to more than one locus), we set the maximum number of loci a read is allowed to map to, to 1 (--outFilterMultimapNmax 1). We also directed STAR to output unmapped reads within the main SAM file by specifying --outSAMunmapped Within, for further downstream analyses. All alignment outputs were sorted by coordinate (--outSAMtype BAM Sort- edByCoordinate) and written to file in standard SAM format. Passed attributes --outSAMattributes (NH HI AS nM NM MD jM jI rB MC) are as defined in the SAM format specifications (Li *et al.* 2009; The SAM/BAM Format Specification Working Group 2023).

**Alignments with STAR**

STARpar = “--runThreadN 16 --genomeDir $genomeDirectory --genomeLoad NoSharedMemory --outSAMtype BAM SortedByCoordinate --outSAMattributes NH HI AS nM NM MD jM jI rB MC --alignEndsType EndToEnd --outSAMunmapped Within --outFilterMultimapNmax 1”

readFiles = “--readFilesCommand gunzip -c --readFilesIn R1.fastq.gz R2.fastq.gz”

$STAR $STARpar $readFiles

STAR+WASP alignment jobs were also run at 8, 16 and 32 threads, maintaining the majority of the parameters applied for the STAR alignments. However, for STAR+WASP runs, our re-implementation of the original WASP algorithm (Van De Geijn *et al.* 2015) was applied and activated with the --waspOutputMode SAMtag. This implementation adds a vW tag to the SAM output where vW:i:1 represents alignments that passed WASP filtering while all other values (vW:i:2-7) represent alignments that did not pass WASP filtering. A description of all vW tags is provided in **Supplementary Table 1**. In addition to the SAM output attributes was the vA, vG and vW attributes which respectively output and specify the variant allele, the genomic coordinate of the variant overlapped by the read, and a flag specifying whether the alignment passed WASP filtering or not. Prior to conducting WASP filtering, we created a directory containing compressed text-based SNP files per chromosome. Each file contains three columns (position, ref allele, alt allele) delimited by space, with the file names containing the name of the respective chromosome. We also supplied VCF files with heterozygous sites on which WASP filtering was done.

**Alignments with STAR+WASP**

STARpar = “--runThreadN 16 --genomeDir $genomeDirectory --genomeLoad NoSharedMemory --outSAMtype BAM SortedByCoordinate --outSAMattributes NH HI AS nM NM MD jM jI rB MC vA vG vW --waspOutputMode SAMtag --alignEndsType EndToEnd --outSAMunmapped Within --outFilterMultimapNmax 1”

readFiles = “--readFilesCommand gunzip -c --readFilesIn R1.fastq.gz R2.fastq.gz”

hetVcf = “--varVCFfile vcf.snv1het”

$STAR $STARpar $readFiles $hetVcf

Similar to STAR and STAR+WASP alignments, running WASP was conducted at 8, 16 and 32 threads for all sample alignments. We followed WASP’s mappability filtering pipeline and used our previously generated text-based SNP files containing genetic variants (SNPs) as inputs. Furthermore, all reads were mapped using STAR, while applying the same parameters used in the above STAR workflow. Next, we identified reads with a likelihood of having mapping biases using WASP’s “find intersecting snps.py” tool. WASP’s operation to correct for allelic mapping biases takes each read that overlaps a SNP; the allele that is present in the read is flipped to match the SNP’s other allele and the read is remapped. A FASTQ file containing reads with flipped alleles is written out for remapping. Reads were remapped using the same approach (STAR) used in the initial alignment phase. Finally, reads where one or more of the allelic versions of the reads failed to map back to the same location as the original read were filtered out.

**Alignments with WASP**

STARpar = ”--runThreadN 16 --genomeDir $genomeDirectory --genomeLoad NoSharedMemory --outSAMtype BAM SortedByCoordinate --outSAMattributes NH HI AS nM NM MD jM jI --alignEndsType EndToEnd --outSAMunmapped Within --outFilterMultimapNmax 1”

readFiles = ”--readFilesCommand gunzip -c --readFilesIn R1.fastq.gz R2.fastq.gz”

hetVcf = ”--varVCFfile vcf.snv1het”

$STAR $STARpar $readFiles $hetVcf

mv $WASPdir/Aligned.sortedByCoord.out.bam $WASPdir/A_sorted.bam

samtools index $WASPdir/A_sorted.bam $WASPdir/A_sorted.bai

$PYTHON $WASP/mapping/find_intersecting_snps.py --is_paired_end --is_sorted --snp_dir $vcfFileDir/ssd_input_snp_dir/SNPdir --output_dir ./ A_sorted.bam

$STAR $STARpar $hetVcf --readFilesCommand gunzip -c --readFilesIn A_sorted.remap.fq1.gz A_sorted.remap.fq2.gz

mv Aligned.sortedByCoord.out.bam Aligned.out.bam

samtools index Aligned.out.bam Aligned.out.bai

$PYTHON $WASP/mapping/filter_remapped_reads.py A_sorted.to.remap.bam Aligned.out.bam A_sorted.keep.bam

All STAR, STAR+WASP and WASP alignments were conducted at full server capacity where each alignment utilized the entirety of all computing resources with no other processes running in parallel. Speed and memory benchmarks for all alignments were measured on a standalone server with a 2 32-core AMD Ryzen Threadripper PRO 3975WX processor running at 3.5GHz, and 528GB of memory (RAM). Resident (built-in) bash commands were used to extract the wall clock and memory usage of each run at the various thread values. Besides conducting read alignments for each sample in a controlled computing environment, using the same workflows as described above for STAR, STAR+WASP and WASP, we conducted similar runs in a shared high performance computing environment, with cluster-based parallel processing. For each sample, we conducted 3 runs per alignment and thread, and averaged benchmarking results across all runs. All benchmarking tests conducted in the shared computing environment were run on a Dell PowerEdge system comprised of Intel Xeon Gold/Platinum (Cascade Lake-SP) processors, 46 compute nodes, 4576 cores (quad Xeon 8260 processors running at 2.4GHz with 24 cores per processor, and 3TB of memory), 56 GPUs (with dual Xeon 6248 processors running at 2.5GHz and with 80 cores each, 768GB of memory, and four Nvidia Tesla V100 GPUs).

Downstream workflows to analyze alignments were conducted using Python (Version 3.9.13) and R (Version 4.2.2) (Code available on [GitHub](https://github.com/rasiimwe/STAR-WASP-reduces-reference-bias-in-the-allele-specific-mapping-of-RNA-seq-reads)). For purposes of plotting and visualization, samples Nucleus nonPolyA R1, Nucleus nonPolyA R2, Nucleus PolyA R1, Nucleus PolyA R2, PolyA R1, PolyA R2, Total R1, Total R2 are all derived from sample NA12878 and originally named NA12878 Nucleus nonPolyA, NA12878 Nucleus nonPolyA Rep, NA12878 Nucleus PolyA, NA12878 Nucleus PolyA Rep, NA12878 PolyA, NA12878 PolyA Rep, NA12878 Total, NA1- 2878 Total Rep respectively.
