## Supplementary Figures for "STAR+WASP reduces reference bias in the allele-specific mapping of RNA-seq reads"

Supplementary Figures 1-8 and Supplementary Table 1 for

“STAR+WASP reduces reference bias in the allele-specific mapping of RNA-seq reads.”

Rebecca Asiimwe, Alexander Dobin

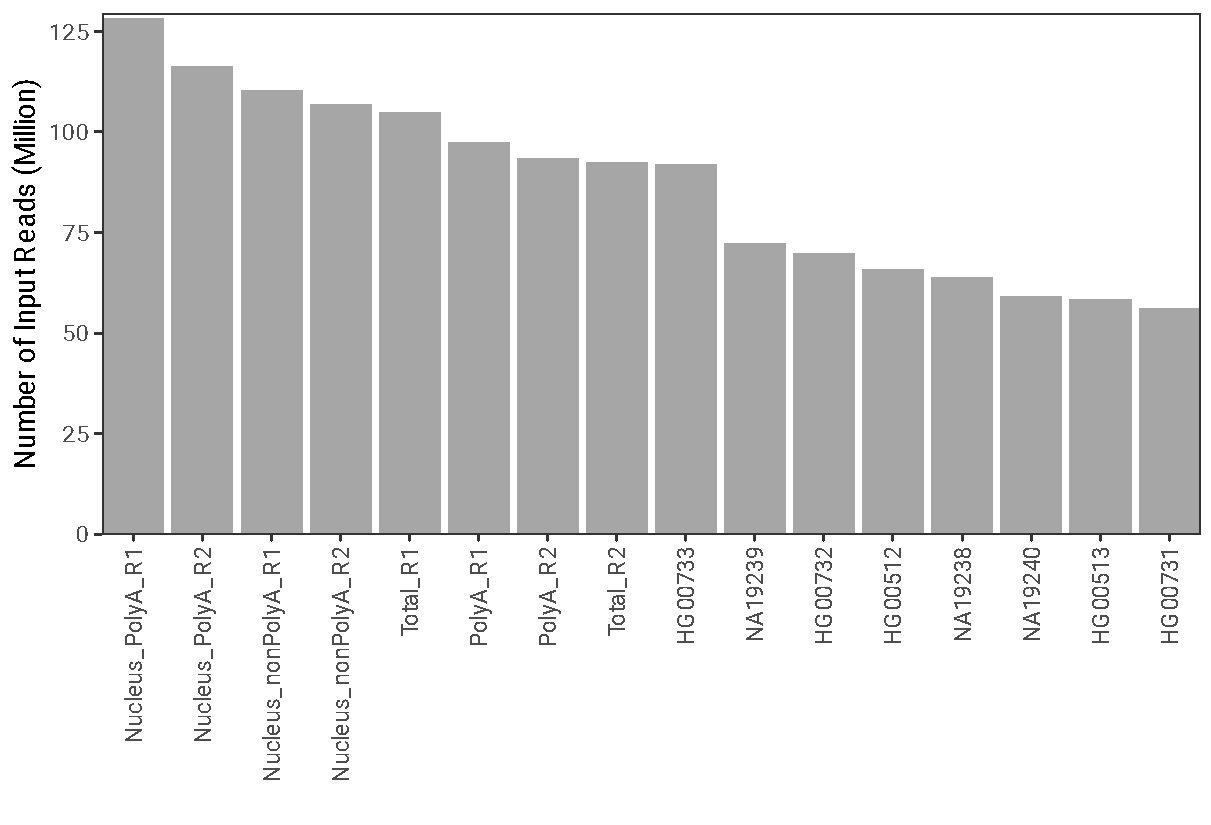

**Supplementary Figure 1: Input reads per sample**. The number of input reads (in millions) for each sample used in this study.

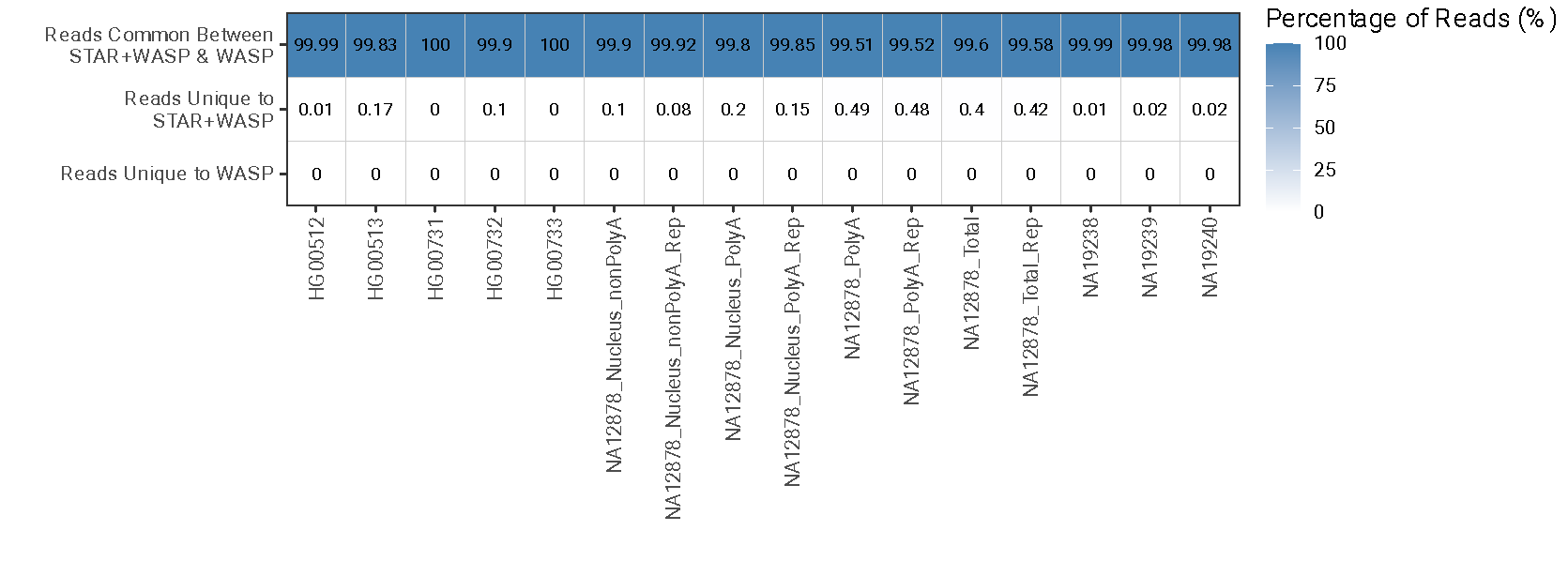

**Supplementary Figure 2:** **Distribution of read percentages for reads overlapping variants discovered by STAR+WASP and WASP**. The top row shows the percent distribution of reads that overlapped variants and were discovered by both STAR+WASP and WASP. The second row shows the percent distribution of reads that overlapped variants and were uniquely discovered by STAR+WASP and not WASP. The third row shows the percent distribution of reads that overlapped variants and were uniquely discovered by WASP (see **Supplementary Table 2**).

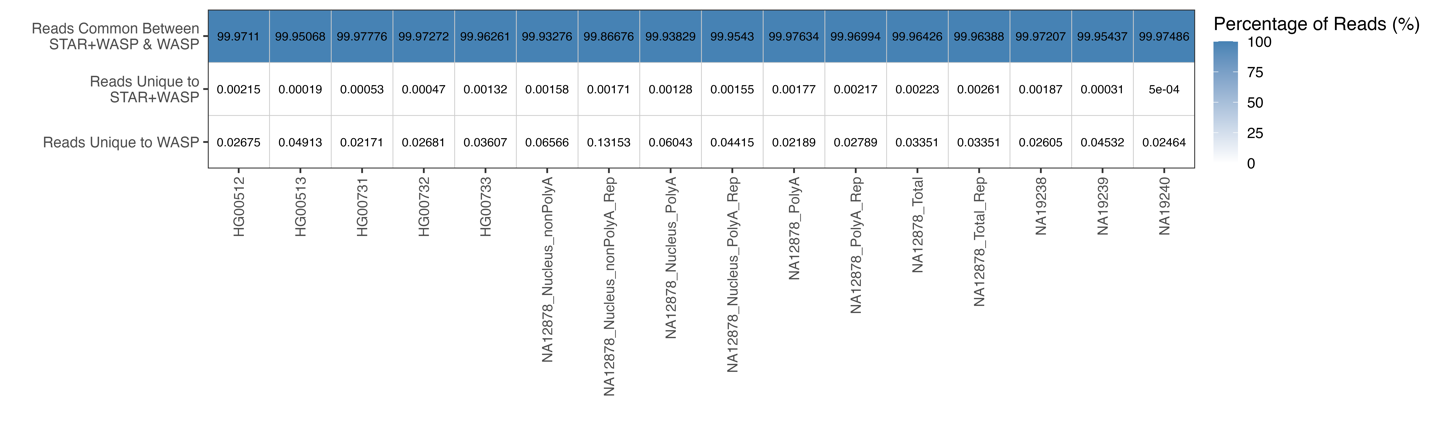

**Supplementary Figure 3:** The percentages of reads that passed filtering (see **Supplementary Table 2**) out of all the reads discovered to overlap variants by both STAR+WASP and WASP. The top row shows the percentage of reads that passed filtering and are common between STAR+WASP and WASP. The middle row shows the percentage of reads that passed filtering and are unique to STAR+WASP. The bottom row shows the percentage of reads that passed filtering and are unique to WASP.

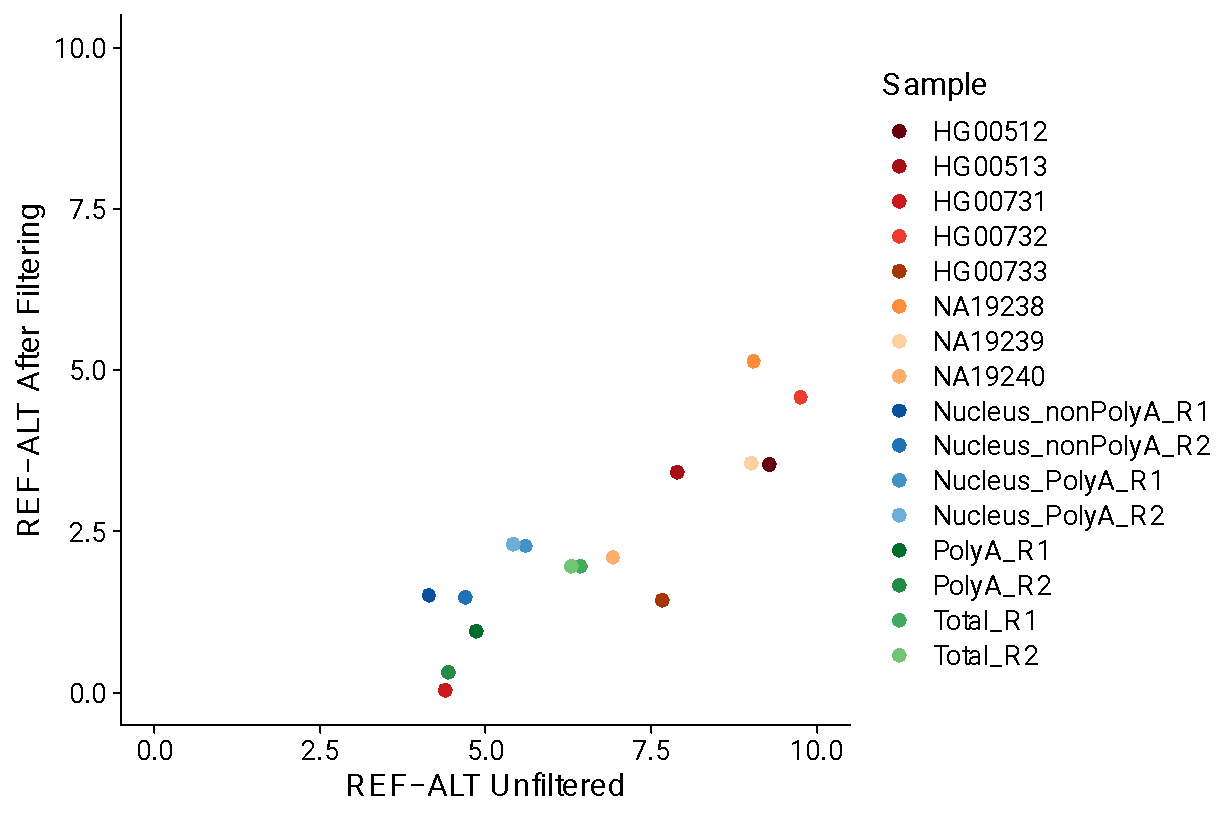

**Supplementary Figure 4:** **Reference bias REF-ALT for before (X-axis) vs. after STAR+WASP filtering (Y-axis)**. REF-ALT is the difference between the percentages of reads that align to the REF allele and those that align to the ALT allele averaged over all variants in each sample.

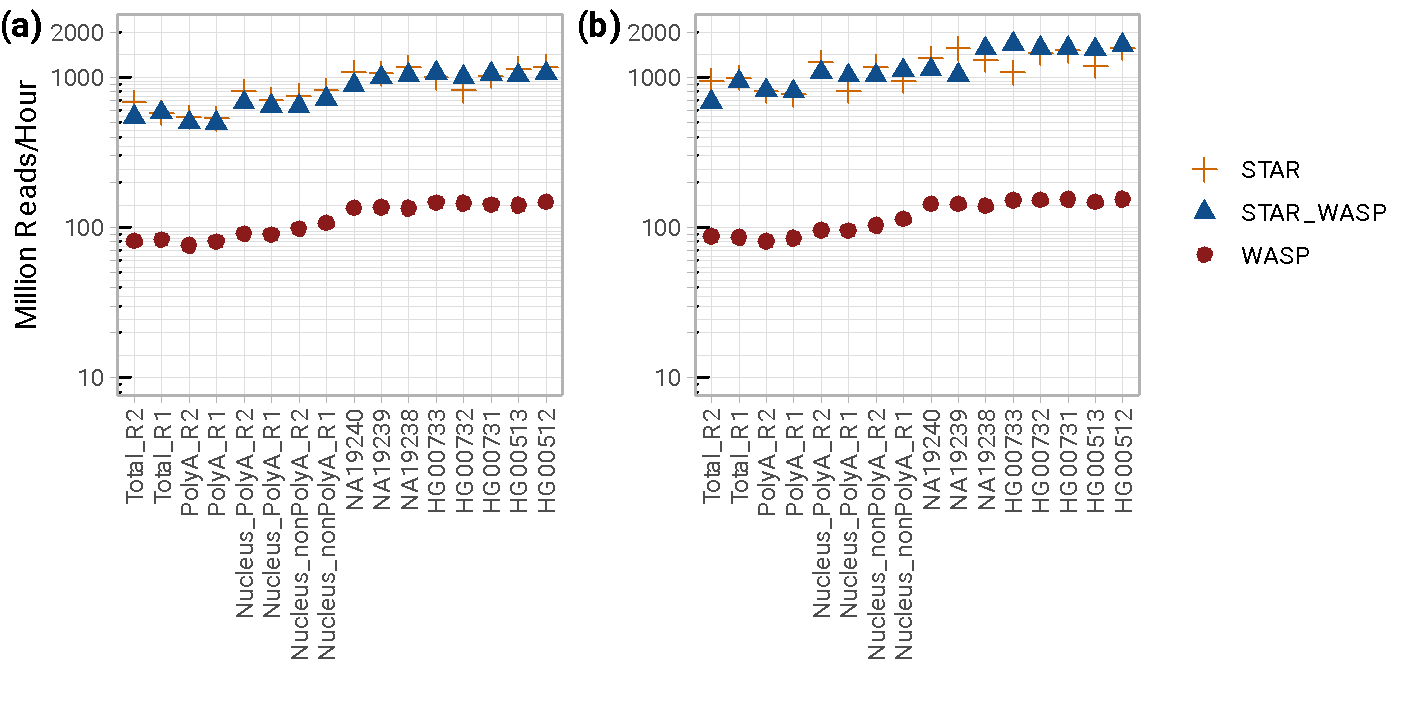

**Supplementary Figure 5: Mapping speed for** STAR, STAR+WASP and WASP for each sample (see **Supplementary Table 3**). Each point represents the overall run time for each sample, colored by the alignment tool. **(a)** 16-thread runs; **(b)** 32-thread runs.

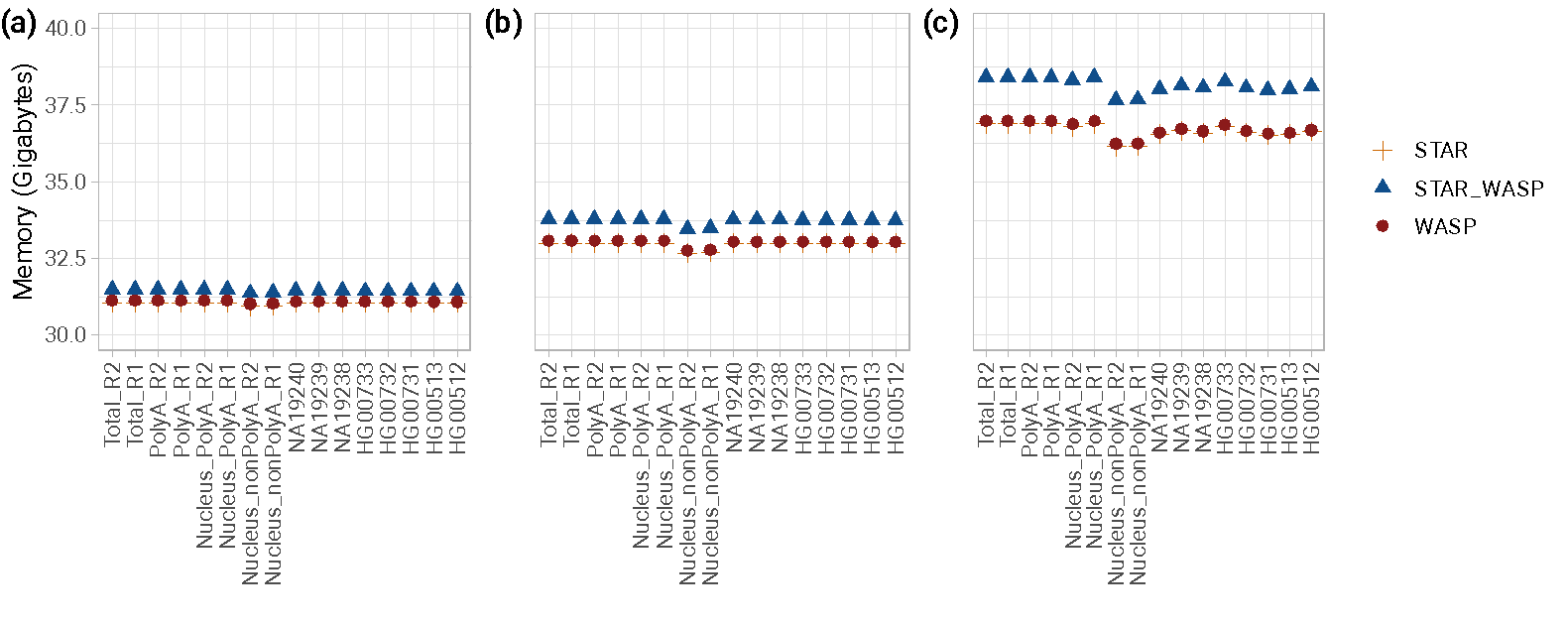

**Supplementary Figure 6: Maximum random access memory (RAM)** used by STAR, STAR+WASP, and WASP for individual sample alignments (see **Supplementary Table 3**). Each point represents the maximum memory used for the alignment of each sample, colored by the alignment tool. **(a)** 8-thread runs; **(b)** 16-thread runs;**(c)** 32-thread runs.

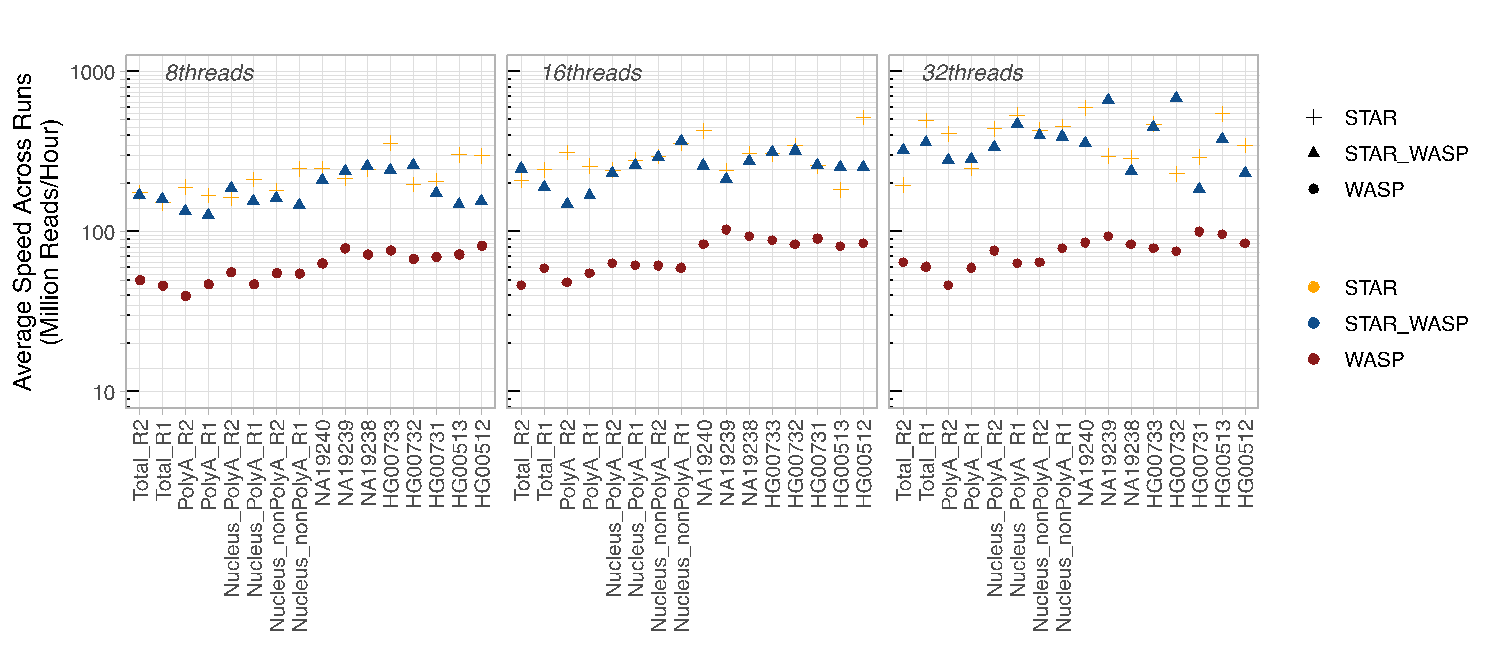

**Supplementary Figure 7: Mapping speed averaged across runs conducted in a shared computing environment** (see **Supplementary Table 4**)**.** Each point represents the total run time averaged across 3 mapping runs conducted on paired-end reads from each sample, by each alignment tool for 8, 16, and 32 threads. Threads are distinguished by plot facet, while alignment tools are distinguished by point shape and color.

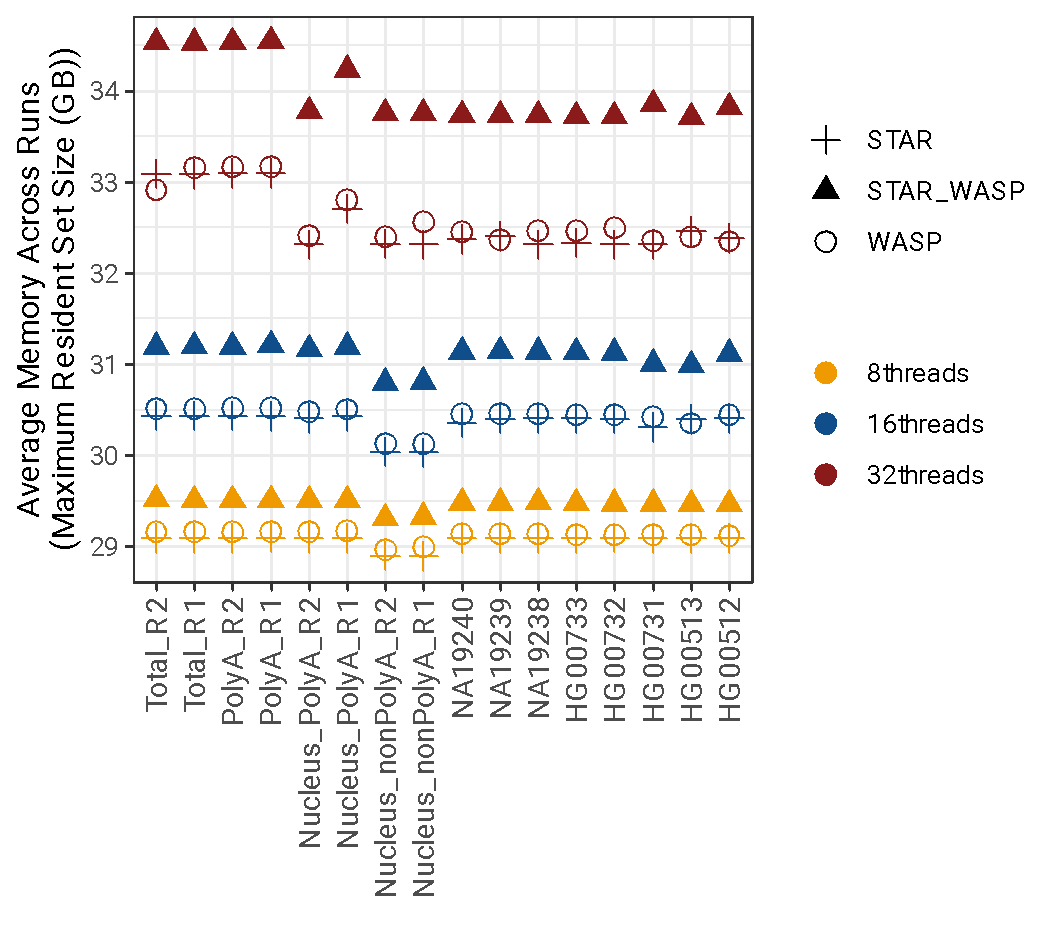

**Supplementary Figure 8: Average maximum memory usage across runs conducted in a shared computing environment** (see **Supplementary Table 4**)**.** Each point represents the average memory used to align paired-end reads from each sample, and by each alignment tool for 8, 16, and 32 threads. The number of threads is distinguished by color, while the alignment tools are distinguished by point shape.

**Supplementary Table 1:** vW Tag Definitions

| **vW Tag** | **Definition** |
| --- | --- |
| vW:i:1 | Alignment passed WASP filtering |
| vW:i:2 | Multi-mapping read |
| vW:i:3 | Variant base in the read is N (non-ACGT) |
| vW:i:4 | Remapped read did not map |
| vW:i:5 | Remapped read multi-maps |
| vW:i:6 | Remapped read maps to a different locus |
| vW:i:7 | Read overlaps too many variants |
